# Mutation load dynamics during environmentally-driven range shifts

**DOI:** 10.1101/333252

**Authors:** Kimberly J. Gilbert, Stephan Peischl, Laurent Excoffier

## Abstract

The fitness of spatially expanding species has been shown to decrease over time and space, but specialist species tracking their changing environment and shifting their range accordingly have been little studied. We use individual-based simulations and analytical modeling to compare the impact of range expansions and range shifts on genetic diversity and fitness loss, as well as the ability to recover fitness after either a shift or expansion. We find that the speed of a shift has a strong impact on fitness evolution. Fastest shifts show the strongest fitness loss per generation, but intermediate shift speeds lead to the strongest fitness loss per geographic distance. Range shifting species lose fitness more slowly through time than expanding species, however, their fitness compared at equivalent geographic distances spread can be considerably lower. These counter-intuitive results arise from the combination of time over which selection acts and mutations enter the system. Range shifts also exhibit reduced fitness recovery after a geographic shift and may result in extinction, whereas range expansions can persist from the core of the species range. The complexity of range expansions and range shifts highlights the potential for severe consequences of environmental change on species survival.

**Author Summary:** As environments change through time across the globe, species must adapt or relocate to survive. Specialized species must track the specific moving environments to which they are adapted, as compared to generalists which can spread widely. During colonization of new habitat, individuals can accumulate deleterious alleles through repeated bottlenecks. We show through simulation and analytic modeling that the process by which these alleles accumulate changes depending upon the speed at which populations spread over a landscape. This is due to the increased efficacy of selection against deleterious variants at slow speeds of range shifts and decreased input of mutations at faster speeds of range shifts. Under some selective circumstances, shifting of a species range leads to extinction of the entire population. This suggests that the rate of environmental change across the globe will play a large role in the survival of specialist species as compared to more generalist species.

## Introduction

The rate of environmental change experienced by organisms plays a major role in driving evolution and determining species survival. Global climate change is just one example of a force driving environmental change. The rate of climate warming is unprecedented in recent history (Huntley 1991) and is predicted to continue into the future (Loarie *et al.* 2009), threatening the survival of many species (Bellard *et al.* 2012, Davis & Shaw 2001, Jump & Peñuelas 2005, Parmesan & Yohe 2003, Thomas 2010). Regardless of the cause of environmental change, organisms must either adapt or shift their range to find suitable environments, and many species already show evidence of range shifts (Chen *et al.* 2011, Frei *et al*. 2010, Grabherr *et al* 1994, IPCC 2007, Kullman 2002, Lenoir & Svenning 2015, Lloyd & Fastie 2003, Parmesan & Yohe 2003, Peñuelas & Boada 2003, Pinsky *et al.* 2013, Sanz-Elorza *et al.* 2003, Sturm *et al.* 2001, Walther 2003, Walther *et al.* 2002). Surviving a range shift is not as simple as tracking an environmental optimum via sufficient dispersal due to the complex genetic, selective, and demographic processes contributing to fitness loss as populations move over geographic space.

Individuals on expanding fronts are known to accumulate deleterious mutations over time and space, leading to fitness loss (termed expansion load, Peischl *et al.* 2013) that could lead to extirpation of local populations or the extinction of species. Expansion load is the consequence of genetic surfing of deleterious mutations at expanding range fronts (Edmonds *et al.* 2004, Klopfstein *et al*. 2006), where inefficient selection due to small population size prevents the purging of deleterious variants, leading to severe fitness loss. This expansion load creates a gradient of fitness across species ranges, where high fitness individuals persist in the core of the species range and low fitness individuals exist at the edge. Theoretical models of range expansions well predict the accumulation of expansion load (Excoffier *et al*. 2009, Gilbert *et al.* 2017, Hallatschek & Nelson 2010, Peischl *et al*. 2013, 2015, Peischl & Excoffier 2015, Travis *et al.* 2007), and empirical evidence of such load continues to emerge (Bosshard *et al*. 2017, González-Martínez *et al.* 2017, Henn *et al.* 2016, Peischl *et al*. 2018, Willi *et al.* 2018). We expect similar processes to occur during range shifts, however, little work has investigated the fitness consequences of a range shift. The combination of variable speeds of spread over the landscape with the lack of a dense, genetically diverse and high fitness species core is expected to greatly impact the dynamics of expansion load at the expanding front. When spread is fast, populations exhibit smaller population sizes at the front leading to stronger genetic drift and thus greater expansion load. Gilbert *et al*. (2017) showed that when range expansions are slowed by the need to locally adapt, the severity of expansion load is reduced. Other processes that slow expansion are also expected to reduce fitness loss during a range shift, such as Allee effects which require a population to reach a given size before growing and expanding further (Stephens *et al.* 1999). Furthermore, the absence of migration from behind the expanding front is also expected to reduce recovery after a shift.

Here, we investigate the loss of fitness due to expansion load in both range expansions and range shifts to understand the important demographic and genetic differences across these scenarios. We assume that range expansions spread at the limit of individuals’ dispersal abilities, while range shifts spread at a speed determined by the rate of environmental change, maintaining a constant population width which expands at the front and recedes at the rear. We also compare how these different demographic scenarios may lead to different dynamics of population recovery, given that gene flow from the species core is a major factor in recovery for expansions and will be lacking in range shifts. We assess the impact that speed of environmental change has on the severity of fitness loss during a range shift. These results have implications for the persistence of species in the face of global climate change and how various demographic scenarios can lead to different outcomes for species in terms of genetic diversity and population fitness.

## Results

### Range shifts lead to greater fitness loss per distance

#### Soft selection

We compared the evolution of mean fitness at the leading edge of an unconstrained range expansion with range shifts in which the speed of the shift is constrained by extrinsic forces such as environmental change. Importantly, the speed of the unconstrained range expansion sets a limit for the upper speed at which a range shift can successfully track a moving environmental niche. We find that rate of fitness loss per generation is less severe in range shifting species than in expanding species (Fig 1A and 1B, Table S1), but the speed at which the range shifts proceed is a key factor determining the rate of fitness loss per generation. When the speed of the shift is close to the speed of a range expansion (speed *v* = 0.2 demes per generation vs. *v* ≅ 0.26 respectively, Fig 1), expansions and shifts, have similar rates of fitness loss per generation (Fig 1A and 1B). Decreasing the speed of range shifts leads to less fitness loss per generation (Fig 1A and 1B), as expected. Surprisingly however, the rate of fitness loss per unit space is greatest at intermediate speeds of range shifts (Fig 1C and 1D). When mutations are fully additive, the fitness of a range shifting species is lower than that of a range expanding species when compared at the same distance travelled (Fig 1C). With fully recessive mutations, faster shifts and expansions initially experience more fitness loss per deme than slower shifts. This is because recessive mutations can be maintained at higher frequencies under mutation-selection balance prior to a shift or expansion, and strong drift at the expansion front leads to rapid expression of these alleles in the homozygous state even though the average number of deleterious alleles per individual remains constant (Kirkpatrick and Jarne 2001, Peischl & Excoffier 2013). This is reflected in the higher number of fixed deleterious variants at the front when mutations are recessive (Fig S1). Slower shifts avoid this initial rapid increase in homozygosity because drift is less strong but do have a steeper slope of fitness loss per space overall and eventually lose more fitness overall as compared to the fastest shifts (Fig 1D). At the slowest speed of range shifts, our simulations deviate from the analytic model (Fig 1A and 1B) because at these slower speeds migration from behind the front has time to reach the range edge, which is not a factor included in our analytic model.

**Fig 1.**
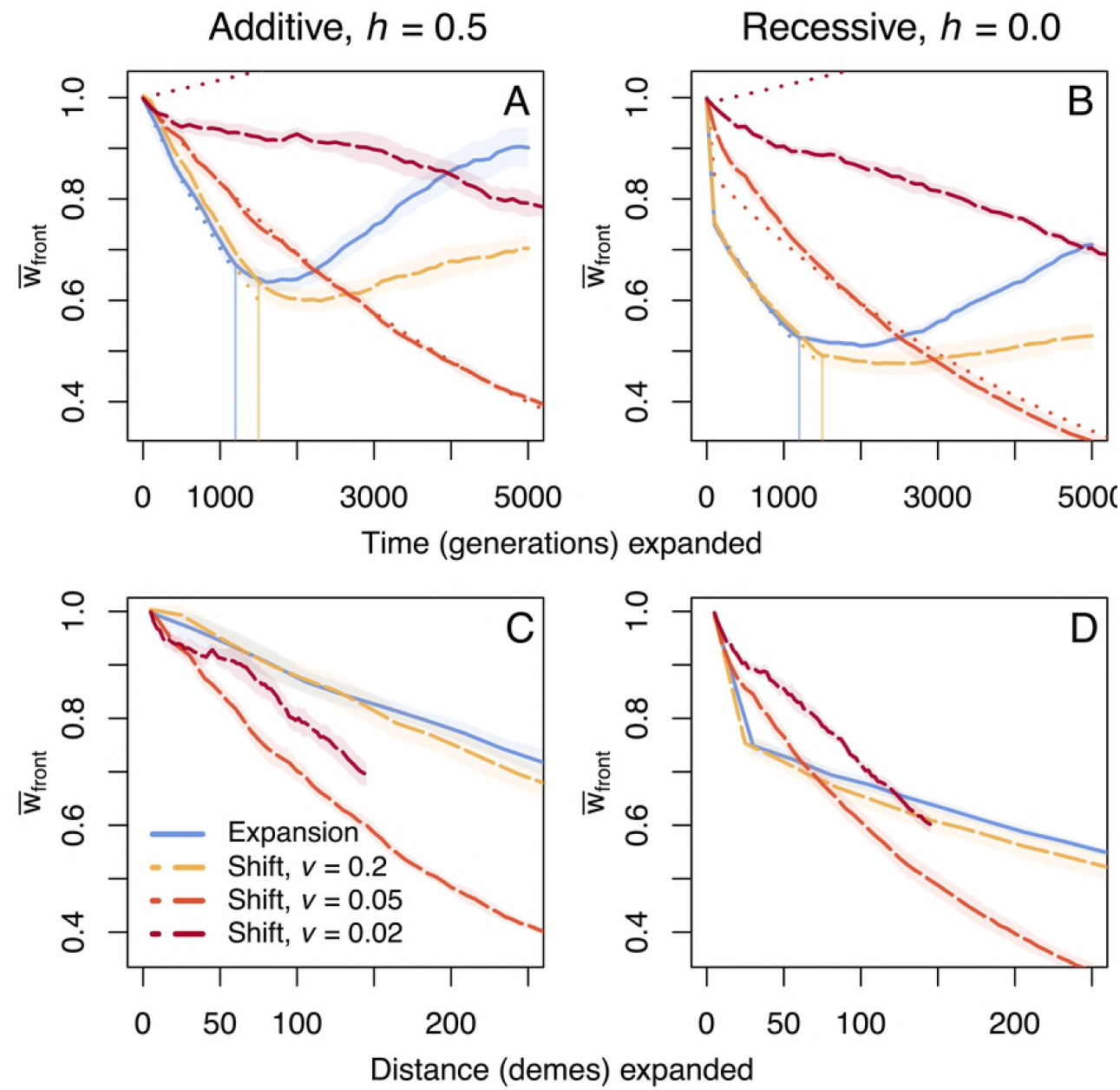
Fitness loss per time and space under soft selection. Trajectories of mean fitness loss over time and space at the expanding front under soft selection show more overall fitness loss for range shifts. Vertical lines indicate when the population reaches the end of the 1×300 deme landscape and expansion is complete. Shaded regions show two standard errors calculated over ten replicate simulations. The fastest shift (*v* = 0.2) expands at a speed closest to the full expansion, and is compared to two slower speed shifts (*v* = 0.05, 0.02). At *v* = 0.02, the population has not crossed the landscape over the time course of the simulation – the end of these lines in C and D are only indicative of this, and not extinction. Analytic solutions for fitness loss over time are shown as dotted lines in panels A and B, where evolution of mean fitness is given by eq. (1) with *F = K m /2*, where *K* is the (diploid) carrying capacity of a deme. The accumulation of fixed deleterious and fixed beneficial mutations for these cases can be seen in Supplemental Fig S1.

To further understand the relationship between the speed of a range shift and fitness loss per unit space, we compared our analytical model to additional simulations (*v* = 0.2, 0.1, 0.066, 0.05, 0.04, 0.033, 0.025, and 0.02 demes per generation; Fig 2). Our model predicts that the fitness loss per unit of space is maximized at a critical speed of approximately *v*_*c*_*≈s* (2*F* − 1) (*2φ* − 1) = 0.056 demes per generation, which matches our simulation with the most severe fitness loss at *v* = 0.05 (Fig 2B). Our model allows us to disentangle the evolutionary forces that govern the accumulation of deleterious mutations during range shifts. As shifts proceed faster, the time taken to colonize a new deme is reduced thereby decreasing the average number of mutations that will spontaneously enter the population (Fig 2C). Furthermore, as shifts proceed faster, population sizes are on average smaller at the front (Hallatschek 2008) leading to more genetic drift and gene surfing. This decrease in *N*_e_ leads to a higher probability of fixation for deleterious alleles and a lower probability of fixation for beneficial alleles (Fig 2D, Peischl *et al*. 2016), resulting in slower range shifts always exhibiting less fitness loss per unit time (Fig 2A). The trade-off between efficacy of selection (more selection during slower shifts) and the amount of influx of harmful mutations during a range shift (more mutations during slower shifts) creates the non-monotonic behavior we find in both the analytic model and simulations (Fig 2B). This non-monotonic behavior persists across a range of carrying capacities and migration rates, with larger population sizes, migration rates or stronger selection leading to faster critical speeds (Supplemental Fig S2). With an increasing influx of deleterious mutations, a wider range of shift speeds lead to greater fitness loss than a range expansion, while increasing the efficacy of selection (either via larger carrying capacities, less severe founder effects, or stronger selection) leads to fewer speeds at which range shifts suffer more fitness loss than expansions.

**Fig 2.**
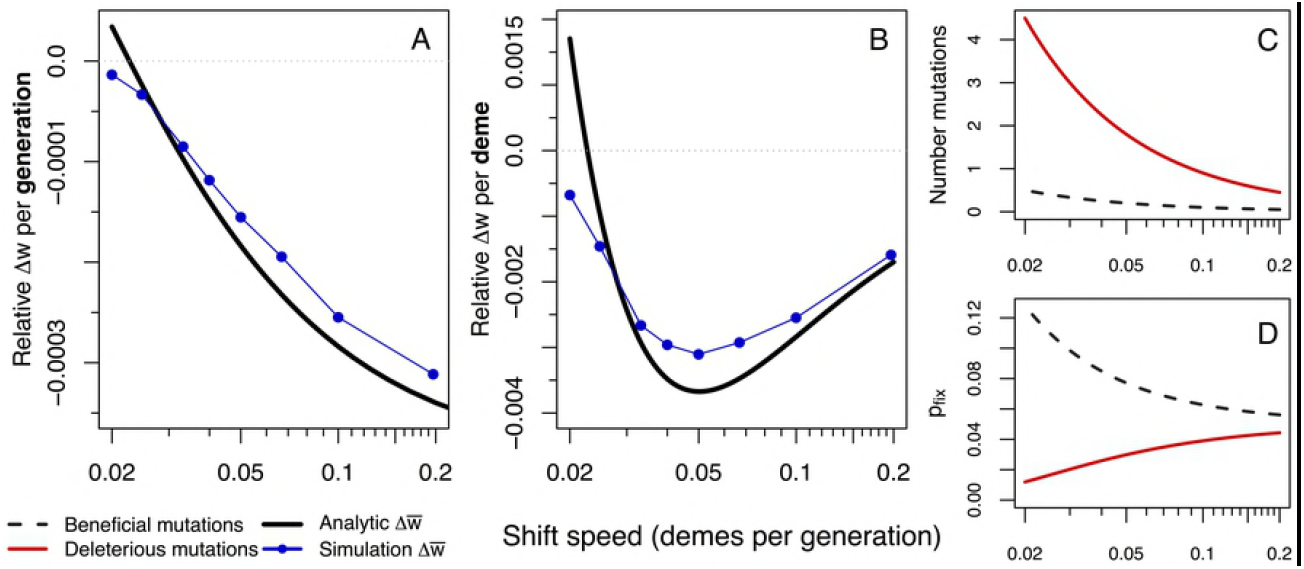
Decomposing fitness loss per time and space. Fitness loss measured per unit time (generations, A) and per unit distance travelled (demes, B). The non-monotonic pattern of fitness loss per distance in B is explained by the combination of mutations entering the population (C) and fixation probability (D) for a given speed of a range shift. Dashed lines indicate beneficial alleles while solid red lines indicate deleterious alleles. The product of fixation probability with number of available mutations produces the fitness change per deme shown in B in solid black. Simulations across speeds are shown in blue, where rates of fitness loss for simulations are calculated within the first 2,000 generations, before beneficial mutations begin to saturate and after generation 100 to ignore initial effects of expansion.

#### Hard Selection

Under hard selection, we find a qualitatively different result where range shifting species can go extinct for the parameter values we used (Fig 3). Because the speed of spread depends on fitness under hard selection, populations can no longer track the speed of environmental change as fitness decreases, resulting in extinction. For the fastest shift (*v* = 0.2), extinction occurs when fitness drops to approximately 0.75-0.78, while the slower shifting species (*v* = 0.05, 0.02) survive longer until fitness decreases to approximately 0.52-0.58. Growth rates are still positive for these fitness values, and stationary populations with this fitness would not go extinct. Our analytical model shows that range shifts can lead to extinction because low-fitness populations can no longer grow sufficiently fast to colonize new habitat, leading to a decline in population size as the landscape disappears behind the shifting range (Fig S3). Range expansions are also slowed due to fitness loss at the expanding front (*v* = 0.176 under the additive model), but extinction does not occur since the population can persist over the whole simulated landscape and be sustained by migrants from the core of the species range. Under the recessive model, fitness loss during expansions is so large that the expanding front stalls until fitness recovers sufficiently to allow further spread. In this case, speed is significantly slowed, and the landscape is not fully crossed during the course of the simulation (populations on average travel 242.6 demes over 5000 generations; *v* = 0.049).

**Fig 3.**
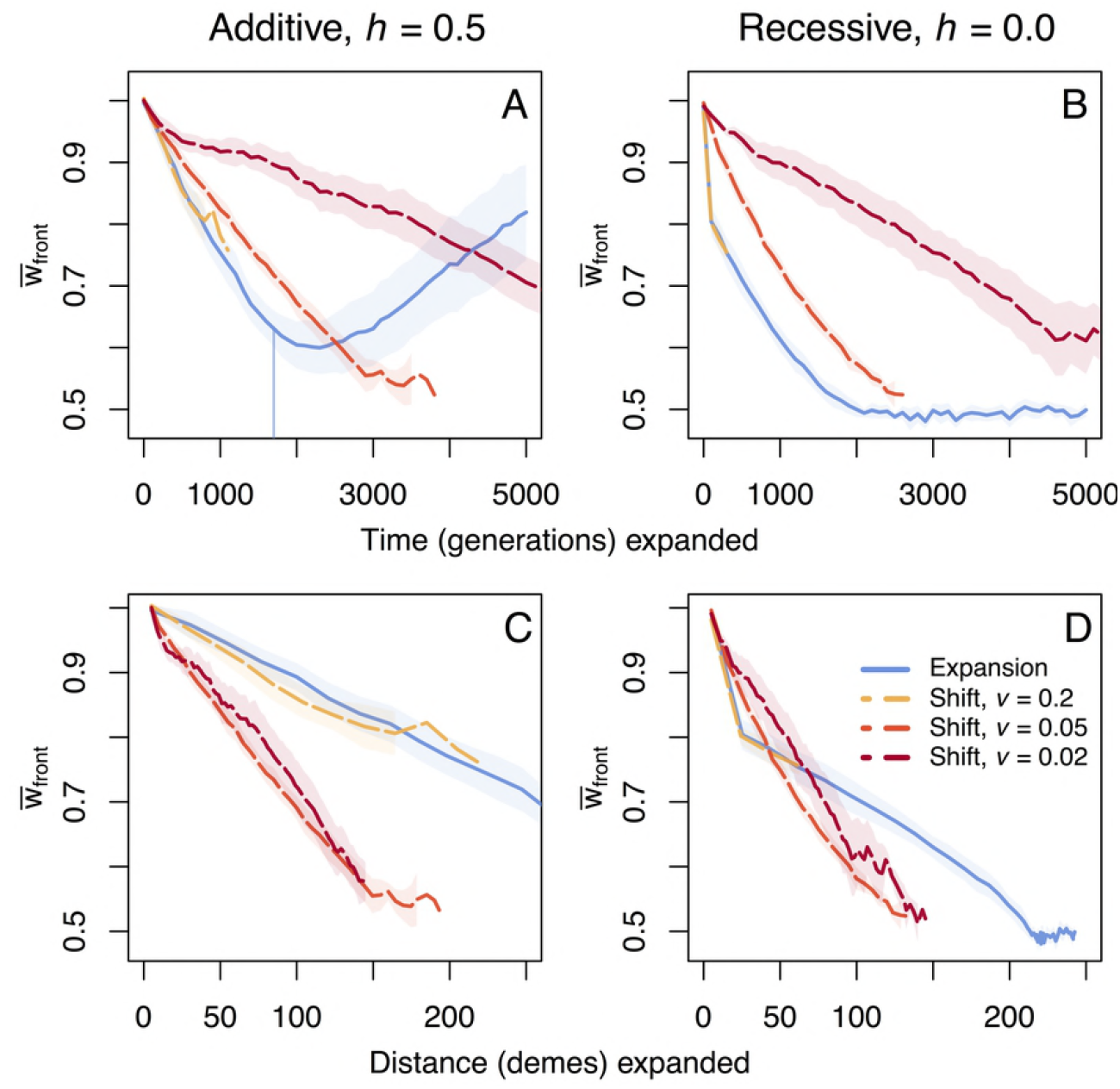
Fitness loss per time and space under hard selection. Trajectories of mean fitness loss over time and space at the expanding front under hard selection during and after range expansions and range shifts. The vertical line in the top left panel indicates when the expansion has reached the end of the 1×300 deme landscape and expansion is complete. This is the only case that finished crossing the landscape during the 5,000 generation time course of simulation, with other cases going extinct or taking more time to spread. Shaded regions show two standard errors calculated over ten replicate simulations.

### Recovery after expansion

In all simulated cases, recovery from accumulated deleterious load is faster and of higher magnitude after a range expansion than after range shifts. Both shifts and expansions exhibit an initial lag in fitness recovery upon crossing the landscape (Fig 1A and 1B) which can be explained by the slower fixation of beneficial mutations once surfing has stopped (Fig S1). Expansions accumulated the least load overall, and thus had less load to recover from (Table S1), yet still show higher rates of recovery than the range shift models (Fig 1A and 1B). Range shifts accumulated more fixed deleterious load than range expansions, and still show minor increases in fixed load after the shift has stopped. In contrast, fixed deleterious load is purged after expansions during this recovery phase (Fig S1). Neutral diversity also returns to a much higher level after an expansion as compared to a shift (average heterozygosity = 0.2 vs. 0.125, respectively; Fig S4). Beneficial mutations show similar rates of increase in fixation during expansions and shifts, but significantly higher rates in the recovery phase for range expansions versus range shifts (Fig S1). Differences in recovery between expansions and shifts arise due to two factors. First, the migration of beneficial variants from the core to the edge of the range reintroduces polymorphism, which is impossible in case of a shift since the core has disappeared. Second, the effective population size is overall much smaller in our range shifts (see Supplemental Figs S5-S6 for further discussion on the effects of *N*_e_ on fitness recovery).

### Incomplete dominance and complex DFEs

We relaxed several assumptions of our mutation model by varying the dominance parameter to include partially recessive mutations and using an exponential distribution of mutational effect sizes (DFE) as described in the Methods. During the initial expansion phase of either shifts or expansions, the rate of fitness loss is minimally affected by these mutational parameters (Figure 4). Only in a single case (*v* = 0.1) does mean fitness loss at the front show a reduced but non-significant rate of fitness loss with an exponential DFE as compared to the additive model with a constant *s* (Fig 4C). Mutational parameters have a stronger impact, however, on the recovery phase after an expansion or shift. When *s* follows an exponential distribution (regardless of the dominance model), fitness recovers at a faster rate as selection increases the frequency of large effect beneficial mutations (Fig S7). The cases with an exponential DFE also show the absence of a lag in fitness recovery once the expansion or shift has stopped. Note that the recovery slows down towards the end of the course of the simulations for range expansions (Fig 4A) because available loci for beneficial alleles begin to saturate (Fig S8B). Importantly, the trade-off modelled between *h* and *s* did not generate results qualitatively different than those obtained for a constant dominance coefficient of *h* = 0.3. This is reassuring, as very little is known about such a trade-off and more research is needed before we can confidently estimate the genomic distribution of dominance coefficients in nature. Thus, while the degree of dominance of new mutations has a bigger impact on fitness loss during the initial expansion phase, the most important factor explaining differences in the rate of recovery in our simulations is the distribution of fitness effects of new mutations.

**Fig 4.**
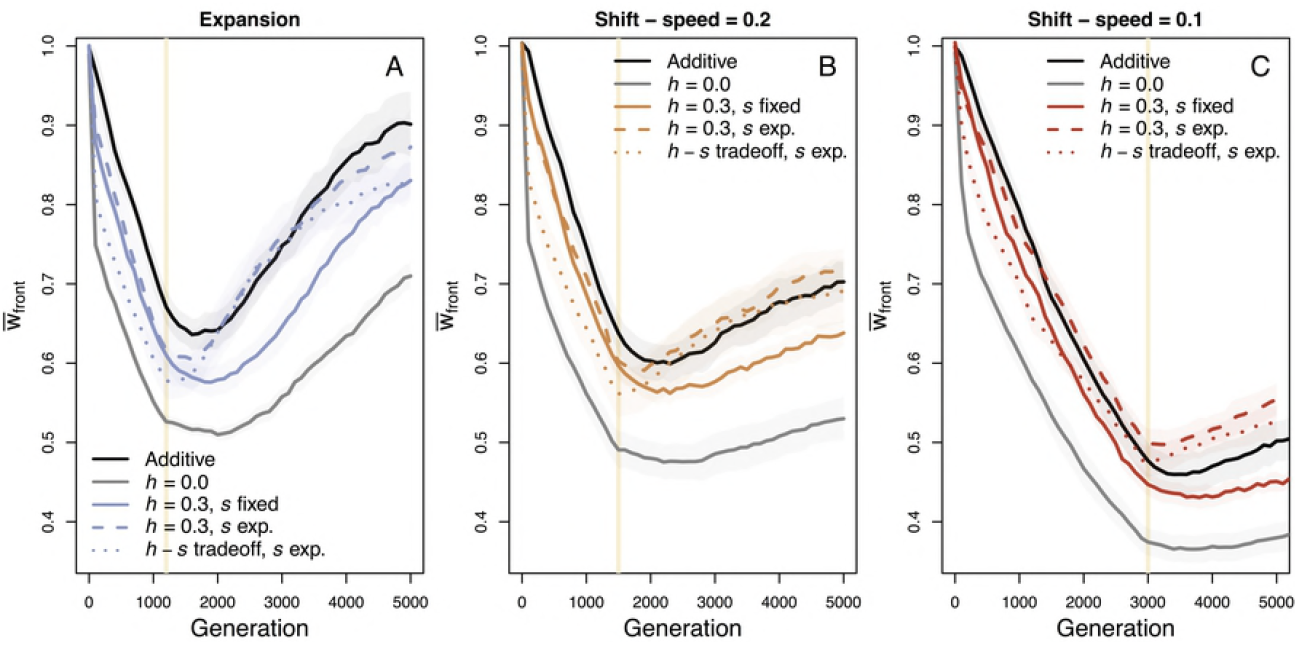
Fitness change over varying mutational assumptions. The assumption of fixed selection coefficients, *s*, and fixed dominance parameters, *h*, are relaxed to compare qualitative outcomes of fitness loss during expansion and fitness recovery after expansion. Shaded regions show two standard errors calculated over ten replicate simulations and the vertical line indicates when the landscape has been crossed and expansion is complete. Our original mutational parameters of fixed *s* and *h* = 0.5 (fully additive) or *h* = 0.0 (fully recessive) are shown in black and gray solid lines, respectively. Colored solid, dashed, and dotted lines show comparison cases of *h* = 0.3 with either constant or exponentially distributed *s* values, or an *h*-*s* trade-off along with an exponential DFE across scenarios of range expansion (A) and our fastest (B) and a slower (C) range shift scenario.

## Discussion

How species modify their ranges in response to environmental change has a large impact on how evolutionary processes unfold within populations. In this study, we have investigated genetic diversity and population fitness both during and after range shifts and contrasted these results to those of a pure range expansion. We uncover two striking results. First, the speed of environmental change driving a range shift is pivotal in determining the dynamics of fitness change over time and space. The severity of fitness loss per unit time qualitatively differs from fitness loss per unit distance, where intermediate speeds accumulate the most expansion load per distance travelled while fastest speeds accumulate the most load per generation time. Second, the mechanism of selection – hard selection or soft selection – leads to qualitatively different outcomes, where range shifts can lead to species extinction under hard selection. These results are vital for predicting population persistence or for implementing reintroduction or other conservation efforts to augment natural populations.

### Fitness loss in time versus space

We have found that since range shifts are forced to proceed more slowly than pure range expansions, fitness loss per unit time is decreased. This is in agreement with previous models of range expansions where it is now well established that faster expansions lead to stronger genetic drift and greater accumulation of deleterious expansion load at the front (Gilbert *et al.* 2017, Hallatschek & Nelson 2008, 2010, Peischl *et al.* 2013). When measuring fitness loss per unit distance travelled, however, we find that range shifts can experience greater fitness loss than expansions for equivalent distances spread. The most severe fitness loss for range shifts is at intermediate speeds, creating a non-monotonic relationship between fitness loss per distance and speed of range shift. This unexpected and counterintuitive pattern of fitness loss results from the fact that the number of generations necessary to travel a given distance determines the number of mutations entering the population as well as the time over which selection may act on those mutations. This effect is seen because the speed at which a range shifting species moves through space is not dispersal- or growth-limited but is limited by the environmental niche which the species occupies. Eventually a range shift (or expansion) that proceeds sufficiently slowly would accumulate no expansion load at the front. Our analytic model (Fig 2) predicts this speed at ≈ 0.0216 demes per generation, while simulations exhibit a slightly slower speed of 0.017 demes per generation (*v* = 1/60) under the additive mutation model and 0.012 demes per generation (*v* = 1/84) for the recessive model (Supplemental Fig S10).

The variable effect of speed on fitness lost during range shifts has important evolutionary implications. The rate of climate change or of anthropogenic changes to the environment will play a major role in determining how fast species must move and thus how much they may suffer from expansion load. Our simulated speed of range shifts is enforced by the environment, meaning that specialist species which must track shifting environmental optima may, under certain conditions, fare better against the input of mutational load when shifts proceed over fewer generations, but only up to the point where too rapid environmental change results in extinction. This may initially bode well for species living on elevational gradients, where environmental change is often greater over shorter distances than latitudinal gradients, requiring less distance travelled to track a moving optimal habitat (until habitat disappears at mountaintops). It is difficult to project our simulated speeds onto real-world speeds of environmental change, as they are specific to our parameter set. Life history traits, generation times, and dispersal abilities of specific species will vary and lead to different degrees of fitness loss for range shifting species. Even though the slowest environmental change is favorable for species survival during range shifts and should imply minimal fitness loss both per time and distance travelled (Fig 2A and 2B), there is clearly no universal optimal speed at which a range shift can proceed, emphasizing the need for species-specific conservation efforts and improved understanding of the interaction between adaptive and dispersive abilities in response to environmental change.

### Hard versus soft selection

At the extreme end of the differences between range shifts and range expansions, we see that range shifting species can go extinct under hard selection, whereas expanding species always survive. Under hard selection population growth depends on fitness. As a consequence, the speed of an expansion is not necessarily dispersal-limited, but instead limited by low fitness and therefore reduced population growth. During range shifts, when fitness drops below the critical level for population sustenance, populations can no longer keep pace with the shifting environment. In the absence of a species core this leads to extinction and is another important effect of the speed of environmental change on the survival of specialist species undergoing range shifts.

Both hard and soft selection are relevant to real-world species and thus to models of range expansions and shifts: organisms that produce offspring in vast amounts may be most subject to local competition and soft selection, while organisms with low reproductive output and high parental investment may experience more hard selection. For example, cane toads, where one mother can produce from 8,000-25,000 eggs in a single clutch (Tyler 1989) would be subject to soft selection and are a classic example of range expansion during their invasive spread throughout northern Australia (Urban *et al.* 2007). On the other hand, many of the world’s large carnivores suffering from human-induced range contractions (Wolf & Ripple 2017) may experience hard selection.

Understanding which species are most likely to undergo range shifts rather than range expansions is thus essential for conserving biodiversity into the future. Specialist species are more likely to shift their range, while generalists are more likely to expand an existing range. Furthermore, specialists that shift over latitudes may travel greater geographic distances than specialists that shift shorter distances over elevation along mountain slopes to track their environment. This may potentially put latitudinally shifting species at greater risk to suffer from expansion load (with the additional caveat that mountainside species will eventually run out of elevation and likely go extinct).

### Demography and mutational parameters impact recovery rates

Recovery from expansion load has not been thoroughly examined in previous studies of range expansions. The presence of a high-fitness species core clearly prevents extinction in the case of hard selection and allows for greater fitness recovery in all cases due to the ability of migrants from behind the expanding front to replenish genetic diversity at the edge. Range shifts lack this recovery mechanism because the core and its high fitness individuals go extinct due to the changing environment. This emphasizes the need to maximally conserve species ranges in their entirety, not only in limited or fragmented sections, and particularly the species range core where individuals are expected to be of higher fitness and possess greater genetic diversity (Eckert *et al.* 2008, Vucetich & Waite 2003).

Effective population size and the connectivity of populations plays a role in recovery from expansion, as is visible in 2-dimensional landscape models (Supplementary Figs S5-S6). Although it is difficult to directly disentangle the effect of the 2-D landscape versus the effect of different effective population sizes, both larger populations and more substructured populations show higher fitness recovery after both expansions and shifts. This is in agreement with previous models which found that 2-D landscapes allow multiple fronts of expansion at which some would experience less fitness loss than others (Peischl *et al.* 2013). Selection can increase the frequency of beneficial mutations and purge deleterious load more efficiently in large populations, and migration among genetically diverse subpopulations with different fixed deleterious alleles can eliminate fixed expansion load. Future simulations implementing even wider 2-D landscapes should be tested, as we would expect shifts to exhibit greater recovery since more genetic diversity would be maintained in a larger population.

The distribution of fitness effects (DFE) of new mutations is also an important factor for population recovery. The true DFE across species and populations still needs to be better understood, but there is general agreement that deleterious mutations have complex and multi-modal distributions (Eyre-Walker & Keightley 2007). Though an exponential DFE did not greatly impact patterns during expansion or shifts in our simulations, post-expansion recovery was greatly improved with an exponentially distributed DFE relative to constant deleterious and beneficial mutational effects (Fig 4), largely because of fixation of highly beneficial variants (Fig S7). The distribution of mutational fitness effects that results after an expansion or shift may also vary depending on the speed of expansion, as has previously been shown by Gilbert *et al.* (2017). Similar to how Balick *et al.* (2015) proposed that the signature left behind by mutations of various dominance levels after bottlenecks could be used to infer the dominance parameter, *h*, experiments measuring fitness recovery after expansions or shifts may provide insight into inferences of the DFE.

### Future Directions

Several interesting future studies are merited from this study. First, further theoretical studies should include the evolution of dispersal. If dispersal rates are able to evolve to higher or lower rates than what is enforced at the start of the simulation, selection may favor less dispersal to reduce expansion load. On the other hand, we would expect range shifting species with higher dispersal abilities to survive longer in the face of environmental change. Burton *et al.* (2010) investigated life history trade-offs in the presence of a dense species core, finding selection for greater dispersal at the edge. However, further investigation is needed to investigate if this result holds in the absence of a dense core. A previous metapopulation model showed higher dispersal evolution as a mechanism of inbreeding avoidance when deleterious mutations are highly recessive (Guillaume & Perrin 2006), emphasizing the importance to better characterize DFEs and dominance parameters along with dispersal evolution to fully understand their effects on expansion load. Second, combining the ability of range shifting species to not only move but also simultaneously adapt to new environmental conditions may lead to qualitatively different results for fitness loss or survival/extinction under hard selection. This type of model could apply to specialist species that may have some adaptive capacity yet still shift to follow their environmental niche.

Last and most important will be to test the predictions of this model with real data. Both experimental evolution and empirical studies in the wild are capable of addressing our results. Bacterial or other experimental studies in the lab could enforce fixed speeds of range shifts and assay fitness across resulting populations. In nature, thorough census data would be necessary to identify species undergoing shifts, but once known comparing the prevalence of deleterious mutations relative to related species that have not undergone range shifts can shed light on these processes. The implications of this study are extremely relevant to biodiversity conservation in today’s world of environmental change, and thus understanding how these factors are realized in real organisms is a vital next step. As climate change proceeds and environments across the globe change at increasingly variable rates, considering the genetic impacts of range shifts may be vital to predict the persistence of many species.

## Methods

We used C++ code for individual-based simulations modified from Peischl & Excoffier (2015; available on GitHub at https://github.com/kjgilbert/ExpLoad) to model range expansions and range shifts over 1- and 2-dimensional discrete space. We follow populations of diploid, monoecious individuals both during the expansion phase as well as after expansion has finished. Random mating occurs within each deme, and generations are discrete and non-overlapping. Dispersal occurs only to adjacent demes with probability *m* = 0.1 per generation and is reflective at the landscape boundaries. Population growth is logistic within each deme (Beverton & Holt 1957). Each deme has a carrying capacity, *K*, of 100 unless otherwise specified and a logistic growth rate model defined by 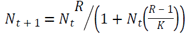, where *R* = 2 and log(*R*) is the intrinsic growth rate. We compare models of both hard and soft selection (Wallace 1975) where carrying capacity and growth rate are constant under soft selection, and carrying capacity and growth rates are proportional to population mean fitness under hard selection (as in Peischl *et al.* 2015).

Both models begin with individuals seeded onto the 5 or 25 left-most demes of a 1×300 or 5×300 landscape grid, for one-dimensional or two-dimensional expansions, respectively, and undergo a burn-in phase of 4,000 generations to reach mutation-selection equilibrium, during which individuals cannot migrate into new, empty demes. In the range expansion model, all empty space on the remaining landscape is opened at the end of the burn-in phase, which allows individuals to colonize and spread at their innate dispersal rate. In the range shift model, both the rate of expansion at the front and the rate of retraction at the rear edge are controlled by maintaining a constant-sized habitat width of 5 or 5×5 demes with *K* > 0, which can be occupied by the population. Range shifts all proceed slower than the range expansions, otherwise they result in extinction. We define a constant speed of range shift as *v=1/T* where *T* is the number of generations between each successive movement forward of the population. *T* = 5 (*v* = 0.2) opens an empty deme at the range front (and forces extinction at the trailing deme) every 5 generations and is our fastest simulated speed of a range shift. This closely approximates the realized speed of the standard range expansion (*v* ≅ 0.25, *T* ≅ 4, Table S1), which results from the maximum growth and dispersal rates used in our model. This model mimics specialist species that must shift their range in either latitude or altitude to track a moving environmental optimum.

Fitness of individuals is determined by 1000 freely recombining, bi-allelic loci and is assumed multiplicative across all loci. We compare both hard and soft selection (see Wallace 1975 for further description of these models). New mutations occur at a genome-wide mutation rate of *U* = 0.1 mutations per diploid individual per generation. Mutations are unidirectional, that is, we prevent back-mutations, and we assume that mutations at 90% of the loci have deleterious fitness effects and 10% have beneficial effects to match previous simulations (Peischl *et al.* 2013, Peischl *et al.* 2015). We ignore beneficial mutations during the burn-in phase, since otherwise all beneficial loci would be fixed for the derived allele before expansion begins and no new beneficial mutations would occur during the expansion. Fitness is scaled to 1 at the end of the burn-in phase to make all scenarios comparable. We examine two main types of dominance models for mutational effects: fixed selection coefficients, *s*, across all mutations of +/- 0.005 (corresponding to a 4*Ks* value of 2) with *h* = 0.5 (additive model) or *h* = 0.0 (fully recessive model), where the fitness contribution at a locus for a heterozygote is 1 + *hs,* and 1 + *s* for amutant homozygote.

In a subset of simulations, we investigate the impact of partial dominance through three additional mutation models: (1) where *h* = 0.3 (partially recessive) across all 900 loci with deleterious effects fixed at *s* = -0.005, (2) *h* = 0.3 and these 900 loci have deleterious fitness effects drawn from an exponential distribution with mean *s* = -0.005, or (3) the same exponential distribution of fitness effects (DFE) for deleterious mutations and a trade-off *h*-*s* relationship. The 100 beneficial loci maintain a constant *h* = 0.5 and have a mirrored exponential distribution to that of the deleterious mutations. More research is needed to understand what distribution of *h* and *s* values is most true in biology, but there is evidence to suggest that more deleterious mutations are more recessive (Manna *et al.* 2011, Agrawal & Whitlock 2011). To test if such a difference in our model affects the outcome, we define an *h-s* relationship with 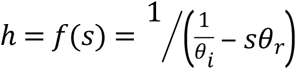, from Huber *et al.* (2017). This relationship is defined by two parameters: we set *θ*_*i*_ = 0.5, which is the intercept of the model defining the value of *h* when *s* = 0, and *θ*_*r*_ is set to 2500 which defines the rate that dominance approaches 0 (fully recessive) as mutation effects become more deleterious (see Supplemental Fig S9). This creates a distribution where dominance approaches complete additivity as neutrality is approached, and dominance approaches complete recessivity as lethality is approached. Even less is known about the DFE of beneficial mutations and hence we model the 100 beneficial loci equivalently across these three comparison cases: an exponential distribution of effect sizes, with mean *s* = 0.005 and a constant *h* = 0.5. To compare levels of neutral genetic diversity post-expansion, 1000 unlinked neutral loci are included in a subset of simulations. To investigate the effects of population substructure and varying effective population size at the expansion front, we also simulated 2-dimensional landscapes, as described in Supplemental Figs S5-S6.

### Analytic model for range expansions and shifts

We compare our simulation results to an analytic model of expansions and shifts under a soft selection model. Peischl *et al*. (2015) showed that the change in mean relative fitness at the front of a linear expansion along an array of discrete demes can be approximated using the following equation:

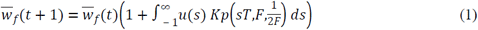

where *u*(*s*) is the mutation rate of mutations with effect *s*, and *P*(*sT,F,P*_0_) = (1 − exp (−*2FsTP*_0_))/(1 − exp (−*2FsT*)) is the fixation probability of mutations with effect *s* and initial frequency *P*_0_ at the front. *F* is the number of founders of a new deme during the expansion, and *T* is the time between two consecutive colonization events. Note that in this model, selection acts during these *T* generations, after which drift acts as a founder effect by randomly sampling *F* individuals. In the case of range expansions, we matched *T* to the average observed speed of range expansion in simulations (*T* = 3.9). We set the relative fitness at the onset of the expansion to 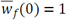 to ensure comparability across results. To compare our results to simulations we assume that *F = K m/2* (Peischl *et al*. 2015).

## Acknowledgements

We would like to thank_______. KJG was supported by EMBO long-term fellowship ALTF2-2016 and LE by Swiss NSF grant No 310030B-166605.

## Data Availability

All simulated data can be regenerated from the parameter sets in Supplementary Table S2. Code for performing the simulations can be downloaded from GitHub at https://github.com/kjgilbert/ExpLoad.

## Supporting Information

**S1 Table. Fitness loss and mutation accumulation across scenarios.** Absolute fitness loss and mutation fixations during expansion per 1-D simulation scenario, averaged over 10 replicate simulations. Cases indicated with a * go extinct before the expansion completes. *T* indicates the number of generations between which the population moves over the landscape and *v* is the speed of spread (inverse of *T*, as defined in the Methods).

**S2 Table. Simulation parameters.** All parameter combinations simulated in the current study to ensure reproducibility of the results. Software code can be downloaded from https://github.com/kjgilbert/ExpLoad. Parameters written in italics within parentheses are the exact software input names used by the simulation. 10 replicate simulations were run with data saved every 100 generations.

**S1 Fig. Mutation fixation through time.** Fixation of deleterious (A, C, E, G) and beneficial (B, D, F, H) mutations at the expanding range front, under soft and hard selection on a 1-dimensional landscape. Vertical lines indicate when the landscape has been crossed and expansion is complete; extinction has occurred for lines that end abruptly. Shaded area indicates two standard errors over 10 replicates.

**S2 Fig. Fitness change through time and space across parameter ranges.** The trade-off between mutations entering the population and selection acting upon these mutations combines to create the non-monotonic pattern of fitness loss seen across speeds of range shifts, as shown by our analytic model. The parameter set in the main text is seen in panels B, E, and H, where carrying capacity, *K* = 100 and migration rate, *m* = 0.1. The impact of beneficial mutations on fitness always decreases with faster speeds due to increasingly inefficient selection (A-C). Deleterious mutations impact fitness non-monotonically across speeds (A-C) because even though more mutations enter the system at slower speeds (more generations pass), selection is more efficient at removing them at slower speeds. Meanwhile at the fastest speeds drift is strongest, but fewer mutations are present (fewer generations for mutational input). D-F show the combined impact of deleterious and beneficial mutations on fitness from A-C. With higher *K* and higher *m*, or extremely low *m*, the non-monotonic pattern of fitness loss per distance travelled is lost. Fitness loss per time (H-I) is always worse at faster speeds.

**S3 Fig. Extinction due to reduced population growth.** As fitness decreases at the front of a range shift due to expansion load (dashed lines), population growth decreases leading to increasingly small population sizes at the expanding front of range shifts (solid lines), under hard selection and with additive mutations. When fitness and thus population size reach a sufficiently low level, the population is no longer able to replace itself as fast as the pace of the shifting environment, resulting in extinction. This occurs more quickly with faster speeds of range shift, since fitness is lost faster through time and populations have less time to recover in size after colonizing new habitat. These analytic approximations qualitatively match our simulations (Figure 3A).

**S4 Fig. Neutral genetic diversity through time.** Neutral diversity over 1000 neutral loci during and after range expansion and shifts at both the expanding edge and in the core (which is calculated as the rear-most deme in range shifts, i.e. the receding edge). Shading indicates 95% confidence intervals over 20 replicates (10 replicates under additive model for selected loci, 10 replicates under recessive model for selected loci). Vertical lines in the left panel indicate when the landscape is crossed and expansion is complete. Slower shifts do not cross the landscape within 5,000 generations. Four various speeds of range shifts are compared.

**S5 Fig. Soft selection 2-dimensional range expansions and shifts.** Range expansions and shifts (*v* = 0.2) in two dimensions are compared for cases where either the population size across the 5-deme-wide front is equivalent to population size in the 1-deme-wide front (2D *K* = 20 vs. 1D *K* =100 and 2D *K* = 100 vs. 1D *K* = 500), or alternatively where the per-deme carrying capacity, *K*, is held constant across comparisons (2D *K* = 100 vs. 1D *K* = 100). Shaded regions show two standard errors calculated over ten replicate simulations. Vertical lines indicate when the landscape has been crossed and expansion is complete

**S6 Fig. Hard selection 2-dimensional range expansions and shifts.** Results for fitness change of 2-D versus 1-D simulations under hard selection. Shaded regions indicate two standard errors over 10 replicates. Vertical lines indicate when the landscape has been crossed and expansion is complete. Absence of a line indicates extinction.

**S7 Fig. Recovery due to beneficial mutations.** Ridgeline plots of allele frequency change through time across the exponential distribution of fitness effect sizes, described in the Methods. Locus allele frequencies have been binned into equal-sized bins of 10 loci each, across the 900 deleterious and 100 beneficial loci, making each line represent 100 bins across the range of the selection coefficient, *s*, rather than 1000 loci. Each individual line across the *y-*axis is a sampled time point, with the start of the simulation being the top- (or back-) most line. Allele frequencies range from 0 to 1 on the *z*-axis.

**S8 Fig. Mutation fixation under various mutation models.** Deleterious (A) and beneficial (B) mutation fixation at the range edge across range expansions and range shifts, over varying mutational models of *h* and *s* as indicated in the figure legend.

**S9 Fig. *h-s* tradeoff.** The *h-s* relationship modelled for deleterious mutations under the *h-s* trade-off scenarios shown in Results Figure 4. (see Methods for description)

**S10 Fig. Equilibrium expansion speeds.** Results under simulations with hard selection for sufficiently slow speeds of range shift show that fitness is on average neither gained or lost at the expanding front, until mutations begin to saturate between generations 2,000 - 3,000. Under an additive mutation model, this speed is realized at 0.017 demes per generation (*v* = 1/60) and under a recessive model at 0.012 demes per generation (*v* = 1/84).

